# STAVelo: deciphering spatiotemporal cellular dynamics via spatial RNA velocity inference from spatial transcriptomics data

**DOI:** 10.64898/2026.10.02.756177

**Authors:** Shang Li, Qunlun Shen, Shihua Zhang

## Abstract

RNA velocity is a powerful tool for deciphering cellular transcriptional dynamics from transcriptomics datasets. However, existing methods have several limitations: some do not model cellular spatial location information, some rely on cell-agnostic constant transcriptional kinetic rates, and some have not been validated for generalization across a wide range of datasets. To this end, we present STAVelo, a graph attention encoder-decoder framework that models spatial information in spatial transcriptomics (ST) data and infers RNA velocity from neural representations of transcriptional kinetic parameters. In quantitative comparisons of multiple datasets from diverse species and platforms, STAVelo outperforms previous methods in capturing cellular migration and development dynamics. In human and mouse brain datasets from different platforms, STAVelo accurately inferred the inside-out migration pattern of neurons across cortical layers. Analysis of mouse kidney and chicken heart datasets captured organ-specific developmental gradients during organ development. Analysis of mouse embryo and placenta datasets with multiple time points uncovered spatiotemporal dynamics from existing states to newly established states. In short, STAVelo provides a spatial view of cellular state transitions and deeper insights into spatiotemporal cellular dynamics in complex tissue structures.

## Introduction

Single-cell RNA-sequencing (scRNA-seq)^1^ captures genome-wide transcriptional profiles at single-cell resolution, facilitating the identification of rare cell populations and reconstruction of cell lineage trajectories^2^. Spatial transcriptomics (ST)^3,4^ retains precise spatial coordinates of gene expression, making up for the limitation of scRNA-seq data lacking spatial location information. To date, ST has been successfully applied to address fundamental biological questions, including depicting tissue structures, characterizing developmental processes, synthesizing three-dimensional tissues and organs, deciphering the tumor microenvironment, and so on ^5–13^.

To elucidate cellular state transitions from static transcriptomics data, researchers have developed transcriptional kinetics models^14^. The core concept is RNA velocity^15^, which quantifies the speed at which cells transition from one state to another, providing a temporal dimension to scRNA-seq analysis. Application of RNA velocity to scRNA-seq data enables investigation of gene expression dynamics at single-cell resolution, prediction of cell trajectories and states, and refinement of trajectory inference through gene regulatory network reconstruction^16–18^. RNA velocity has become an important tool for understanding cell differentiation and development processes in complex biological systems^19,20^.

Current studies demonstrate extensive algorithmic investigations into transcriptional kinetics from scRNA-seq data globally, while studies on ST data remain relatively limited. Pioneering work in this field includes two foundational algorithms: Velocyto^15^ and scVelo^21^. Velocyto is introduced under a steady-state model and estimates RNA velocity across all cells by leveraging deviations in the ratio of unspliced relative to spliced mRNA within transcriptionally stable cells. Although computationally efficient, this method is sensitive to transcriptional noise, affecting velocity estimation. In contrast, scVelo employs a probabilistic dynamical model at its core, iteratively estimating gene-specific kinetic rates and cell-specific latent time via an expectation-maximization algorithm, making it suitable for complex datasets.

Building on prior work, researchers have developed several algorithms to predict RNA velocity for scRNA-seq data. Deep learning-based approaches cellDancer^22^ and DeepVelo^23^ use fully connected and graph convolutional networks to learn kinetic rates and predict RNA velocity, supervised by expression differences among similar cells. Hybrid frameworks that combine dynamical models with deep learning include veloVI^24^ and VeloVAE^25^, which estimate kinetic rates and latent time via expectation maximization or evidence lower bound optimization. Other methods refine kinetic equations: UniTVelo^26^ applies radial basis functions for flexible kinetic modeling, while SDEvelo^27^ adopts stochastic differential equations to capture transcriptional stochasticity and gene-gene crosstalk. Because scRNA-seq data lack spatial location information, the reliability of RNA velocity inferred by these algorithms in real tissue space remains unknown.

In recent years, researchers have developed several algorithms for ST data, such as TopoVelo^28^, CCCvelo^29^, spVelo^30^, and STEER^31^. TopoVelo infers RNA velocity using spatially coupled differential equations and graph neural networks. Yet, its biologically unrealistic assumption of universal splicing and degradation rates hinders large-scale application. CCCvelo jointly estimates RNA velocity and cell communication dynamics constrained by prior ligand-receptor signaling networks, whose performance is susceptible to network incompleteness and annotation errors. spVelo integrates variational autoencoders and graph attention networks, with maximum mean squared error regularization for cross-batch RNA velocity estimation. STEER integrates a spatially informed graph attention autoencoder with a kinetically guided mixture-of-experts to decouple multiple kinetics. spVelo and STEER have only been applied to a small number of ST datasets; their generalization ability is unknown. Continuous advancements in ST technology have generated many high-resolution ST datasets, but the performance of these algorithms on these datasets has not yet been systematically evaluated.

To this end, we have developed STAVelo, a graph attention encoder-decoder deep learning method to infer RNA velocity from ST data. STAVelo first introduces a neural representation of transcriptional kinetic parameters using a spatially-coupled expression encoder and a kinetic decoder. STAVelo then infers RNA velocity using the neural representation of kinetic parameters and expression differences between spatially adjacent cells. We detail the development and validation of STAVelo, demonstrating its superior generalization ability in predicting RNA velocities in ST datasets across multiple species and platforms. Moreover, STAVelo can reveal spatiotemporal developmental patterns on ST datasets with complex spatial gene expression profiles at multiple time points.

## Results

### Overview of STAVelo

The transcriptional kinetics model the mRNA lifecycle from synthesis to degradation, incorporating DNA transcription rate α, unspliced mRNA abundance *u*(*t*), splicing rate β for unspliced mRNA, spliced mRNA abundance *s*(*t*), and degradation rate γ for spliced mRNA (**Fig. 1a**). The process is formally described by the kinetic equations:

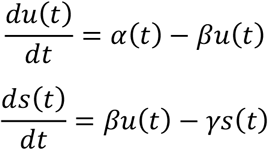

where time *t* denotes biological time (e.g., differentiation stages) rather than physical time. Building upon this, RNA velocity is defined as the instantaneous rate of change 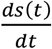 in spliced mRNA abundance at time *t*.

**Figure 1.**
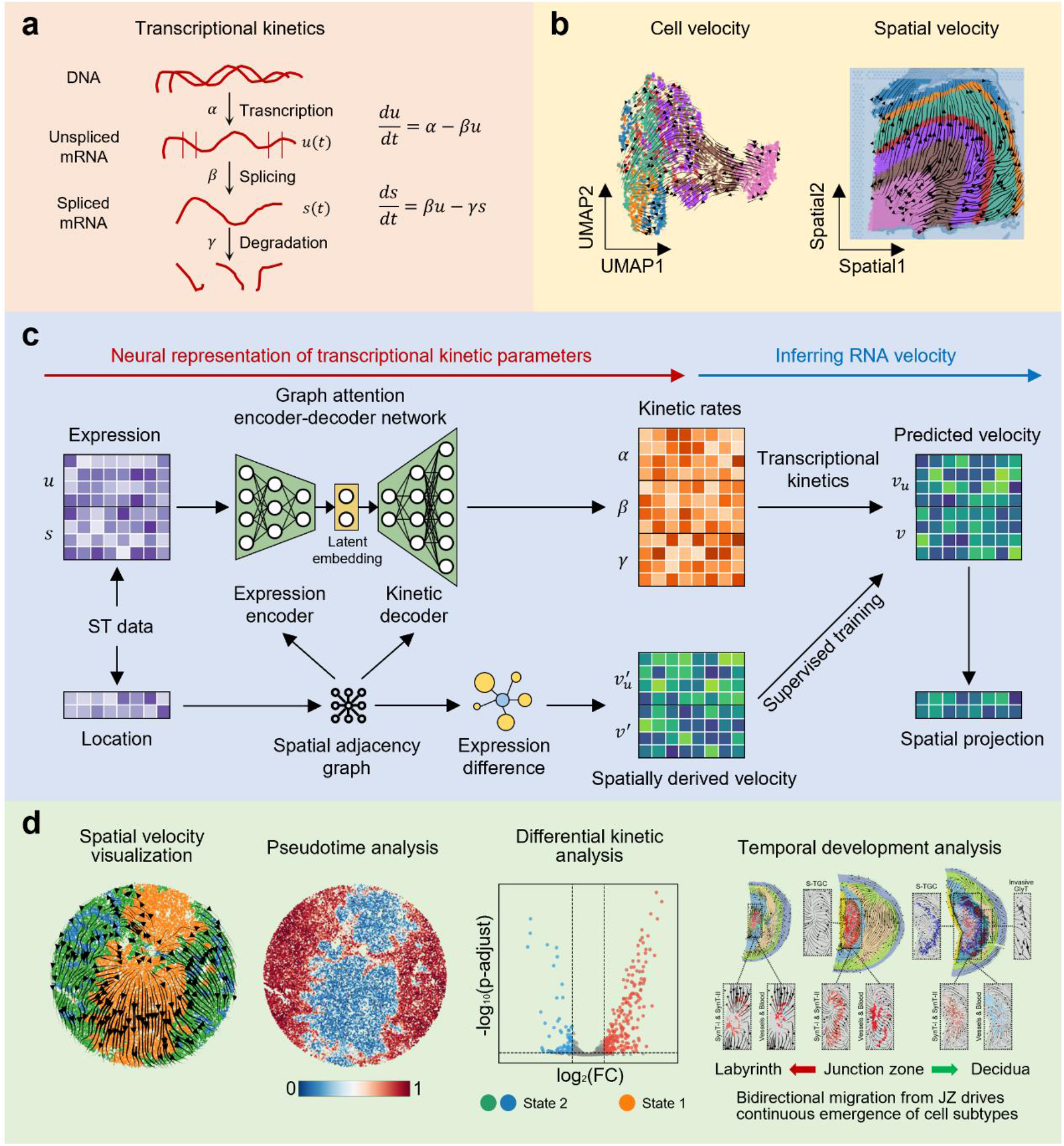
Overview of STAVelo. **a**, Schematic diagram of transcriptional kinetics. The transcriptional kinetic equations model the transcriptional dynamics process for transcription, splicing, and degradation of RNA. **b**, Schematic diagram of cell velocity and spatial velocity. Compared to cell velocity, spatial velocity utilizes spatial locations to reflect spatial development trends. **c**, Model architecture of STAVelo. The graph attention encoder-decoder framework takes the normalized gene expression profiles (unspliced and spliced) and the spatial locations of the ST data as input and outputs the kinetic rates. The predicted velocity is calculated via the transcriptional kinetics model. Spatial locations are used to construct the observed velocity for supervised training. After training, STAVelo can predict RNA velocity of the ST data. **d**, Downstream analysis of STAVelo. STAVelo can be applied to spatial velocity visualization, pseudotime analysis, differential kinetic analysis, and temporal development analysis.

STAVelo employs the above transcription kinetic model (**Fig. 1a**) to infer RNA velocity from ST data. The projection of RNA velocity onto tissue space, termed spatial velocity, exhibits distinct biological significance compared to cellular velocity derived from scRNA-seq data. In scRNA-seq datasets featuring well-defined developmental lineages, cellular velocity reflects a cell*’*s potential to differentiate along specific trajectories. Conversely, ST data often encompasses mature tissues without explicit developmental lineages, instead revealing relative developmental relationships between tissue regions. Consequently, spatial velocity characterizes the relative developmental order among tissue domains (**Fig. 1b**).

STAVelo consists of neural representations of transcriptional kinetics parameters and RNA velocity inference (**Fig. 1c**). STAVelo first adopts a graph attention encoder-decoder to model gene expression and spatial information. Spatial coordinates of spots construct a spatial adjacency graph, where unspliced and spliced expression values per spot serve as node features. The expression data and spatial graph are jointly fed into the expression encoder. Graph attention mechanisms learn inter-spot attention weights, generating low-dimensional latent representations that integrate both gene expression and spatial information. The kinetics decoder subsequently outputs cell-gene-specific transcriptional kinetic rates. STAVelo then calculates the predicted velocity via the transcription kinetics model. The spatially derived velocity of each cell is calculated as the expression difference between the cell and spatially adjacent ones, which supervises neural network training. The trained network ultimately predicts RNA velocity vectors, which can be projected onto tissue space for visualization and further applied for pseudotime analysis, differential kinetic analysis, and temporal development pattern analysis (**Fig. 1d**).

### STAVelo accurately infers the inside-out migration pattern of neurons in human cerebral cortex data

We applied STAVelo to human dorsolateral prefrontal cortex (DLPFC) 10x Visium data^32^. Researchers have manually annotated this dataset with white matter (WM) and six cortical layers, serving as ground truth for evaluation. On section 151676, STAVelo inferred an inside-out migration pattern from WM to the six cortical layers (**Fig. 2a**), consistent with published findings on this pattern and the sequential maturation from layer 6 to layer 2^33,34^. Other methods produced biologically implausible spatial RNA velocity and pseudotime inferences, predicting reversed migration direction (**Fig. 2a**). Quantitative metrics demonstrated STAVelo*’*s superior performance in both spatial CBDir and spatial ICVCoh scores compared to alternative approaches (**Fig. 2b**), confirming its accuracy in predicting cellular migration dynamics. UMAP visualization and PAGA graphs of the kinetic rates revealed clear progression from WM through all six cortical layers (**Fig. 2c**), further evidencing effective spatial information integration via the graph attention network.

**Figure 2.**
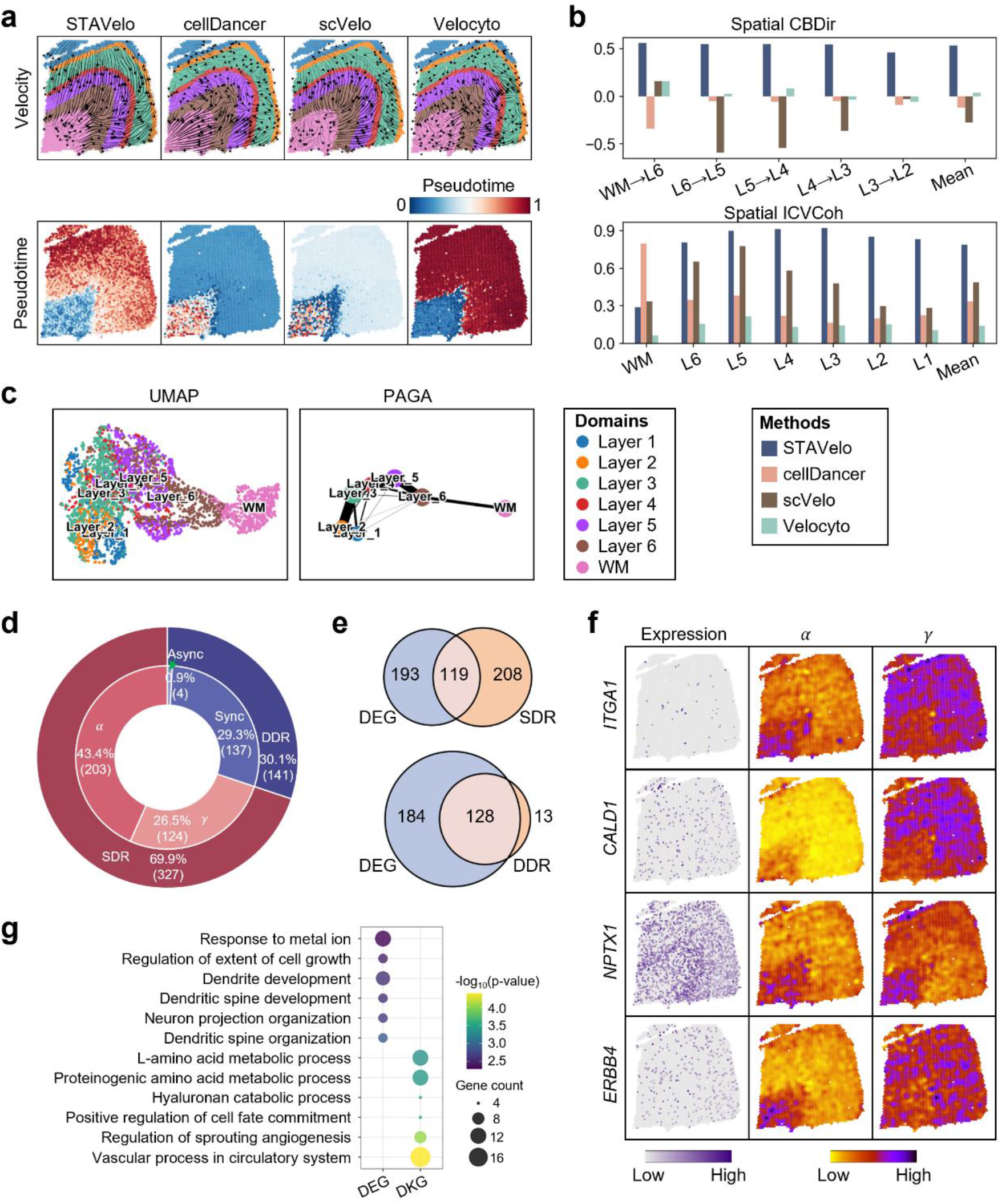
STAVelo infers the inside-out migration pattern of neurons in the human DLPFC section 151676. **a**, Spatial visualization of RNA velocity and pseudotime predicted by STAVelo, cellDancer, scVelo, and Velocyto. Spots are colored by the manually annotated spatial domains. **b**, Bar chart showing spatial CBDir and ICVCoh scores of STAVelo, cellDancer, scVelo, and Velocyto. **c**, UMAP visualization and PAGA graph of kinetic rates. **d**, Double doughnut chart showing the proportion of each category of differential kinetic genes (DKGs), including single differential rate (SDR) and double differential rate (DDR) categories, with DDR further divided into synchronous (Sync) and asynchronous (Async) patterns. **e**, Venn diagrams illustrating overlap between differentially expressed genes (DEGs), SDR, and DDR. **f**, Spatial expression, transcription, and degradation parameter visualization for representative DKGs. **g**, Bubble plot showing enriched GO terms of DEGs and DKGs.

We subsequently evaluated STAVelo and comparator methods on sections 151507 and 151669 (**Supplementary Fig. S1a**). Results from section 151507 showed STAVelo slightly underperformed cellDancer but surpassed both Velocyto and scVelo (**Supplementary Fig. S1b**). Conversely, on section 151669, STAVelo outperformed all methods while cellDancer yielded entirely incorrect results (**Supplementary Fig. S1b**), demonstrating STAVelo*’*s cross-sectional robustness. Ablation studies validating the necessity of the graph attention encoder-decoder network compared STAVelo*’*s predictions against observed velocities (denoted as Obs-velo) (**Supplementary Fig. S2a**). STAVelo achieved higher spatial CBDir scores than Obs-velo in most cases (**Supplementary Fig. S2b**), with correctly oriented spatial RNA velocity vectors. In contrast, Obs-velo produced predominantly erroneous directional inferences (**Supplementary Fig. S2b**), suggesting that denoising via the graph attention encoder-decoder network is essential for accurate spatial RNA velocity estimation.

Similar to STEER, we next identified differential kinetic genes (DKGs) by comparing WM with cortical layers (layers 1 to 6) to dissect kinetic features of state transition from WM to cortical layers. DKGs include only singular differential rate (SDR) and double differential rate (DDR) genes, and the synchronous (Sync) pattern accounts for the vast majority of DDR genes; the asynchronous (Async) pattern is very rare (**Fig. 2d**). Differential expression genes (DEGs) have only a small overlap with SDR genes, but a large overlap with DDR genes (**Fig. 2e**); the combined effects of transcription and degradation lead to significant differences between WM and cortical layers. DKGs, which are not DEGs, contain some important genes that regulate neuronal migration, cortical stratification, and synapse formation, such as *ITGA1, CALD1, NPTX1*, and *ERBB4*. They do not have significant differences in expression, but have significant differences in kinetic parameters (**Fig. 2f**). GO enrichment further supported this concordance: DEGs only capture terms associated with neuronal phenotype that have already matured and whose dendritic synapses are fixed; DKGs specifically capture regulatory events during the intermediate transitional stages of neuronal migration, fate pre-excitation and metabolic reprogramming (**Fig. 2g**). These results also indicate that cell-gene-specific kinetic parameters represented by neural networks help capture kinetic signals related to cell migration and developmental dynamics, which cannot be captured by static transcriptomics data and cannot be achieved by velocity inference methods based on globally uniform kinetic parameters.

### STAVelo accurately infers the developmental relationships of important tissue domains in mouse brain data

We applied STAVelo to mouse brain data, including sagittal-posterior and coronal sections from 10x Visium and a coronal section from Slide-seqV2^35^. We annotated the key brain structures based on STAGATE^5^ clustering results, including the cerebral cortex, hippocampus, thalamus, hypothalamus, fiber tracts, midbrain, cerebellum, and hindbrain. Among all sections, STAVelo revealed an inside-out pattern of cellular migration and maturation in the cerebral cortex (**Fig. 3a**), consistent with the early discovery^33,34^. Furthermore, STAVelo reflected the developmental gradient from the hippocampus to the cerebral cortex among all sections (**Fig. 3a**), confirming that synaptic remodeling in the cerebral cortex ends later than in the hippocampus^36^. For all sections, STAVelo*’*s spatial ICVCoh scores were higher than those of the other competing methods (**Fig. 3b**). Other methods did not correctly infer such developmental relationships (**Supplementary Fig. S3a-c**).

**Figure 3.**
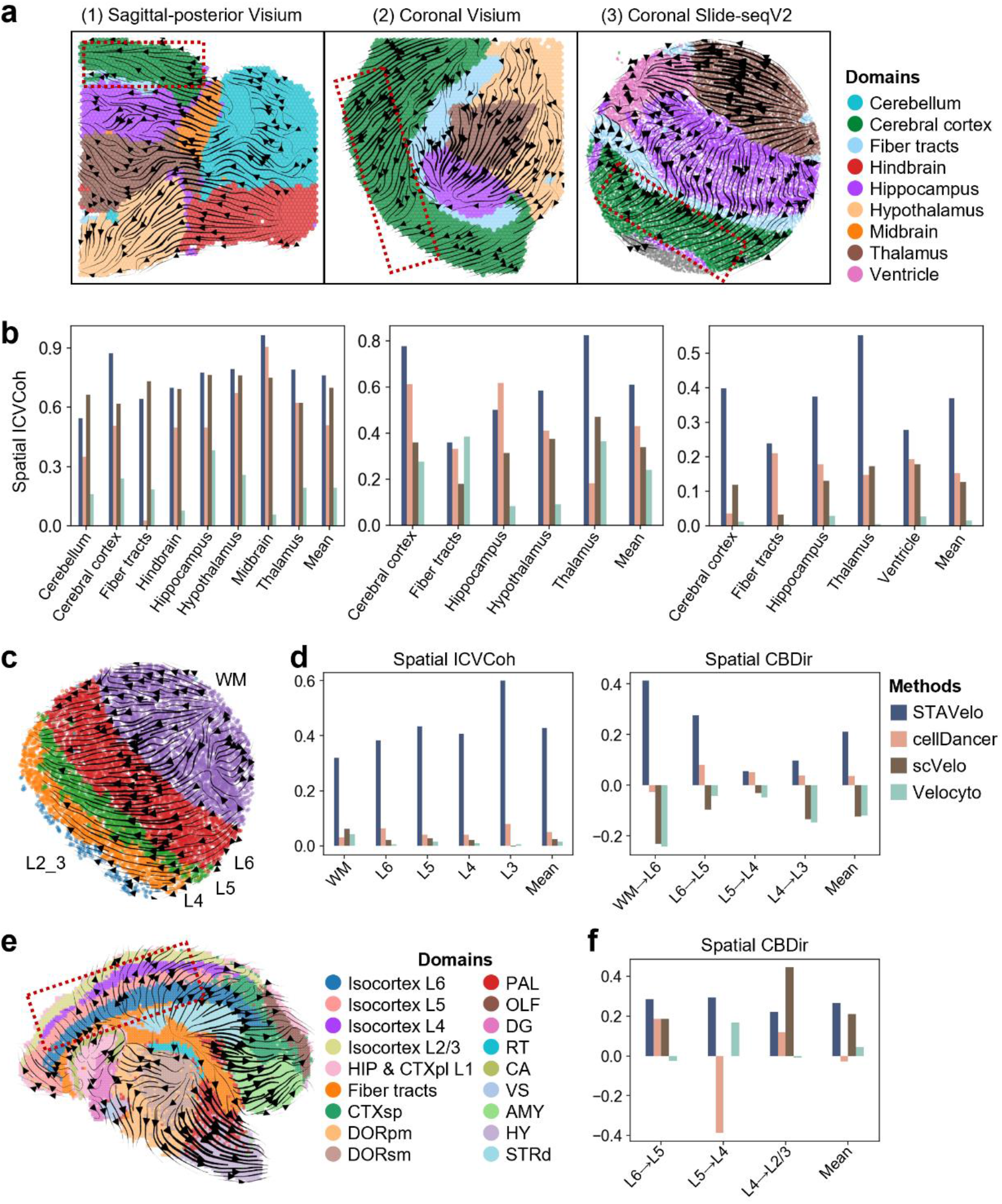
STAVelo infers developmental relationships among important tissue domains in mouse brain data from different sections and platforms. **a**, Spatial visualization of predicted RNA velocity for a sagittal-posterior section and a coronal section from the 10x Visium platform and a coronal section from the Slide-seqV2 platform. Spots/beads are colored by the spatial domains annotated according to the STAGATE clustering results. **b**, Bar chart showing spatial ICVCoh scores of STAVelo, cellDancer, scVelo, and Velocyto for (**a**). **c**, Spatial visualization of predicted RNA velocity for mouse cortex Slide-seqV2 data. Beads are colored by the spatial domains annotated according to the STAGATE clustering results. **d**, Bar chart showing spatial CBDir scores and ICVCoh scores of STAVelo, cellDancer, scVelo, and Velocyto for (**c**). **e**, Spatial visualization of predicted RNA velocity for mouse coronal hemibrain Stereo-seq data. Cells are colored by the spatial domains annotated by the original authors. **f**, Bar chart showing spatial CBDir scores of STAVelo, cellDancer, scVelo, and Velocyto for (**e**).

We also applied STAVelo to mouse cortex Slide-seqV2 data^37^ and mouse coronal hemibrain Stereo-seq data^38^. For mouse cortex Slide-seqV2 data, WM and several cortical layers were annotated based on STAGATE clustering results. STAVelo accurately inferred the cellular migration direction from WM to several cerebral cortices (**Fig. 3c**). STAVelo showed higher spatial CBDir score and spatial ICVCoh score than those of other methods (**Fig. 3d**). Among them, cellDancer had a higher CBDir score between some cerebral cortices than STAVelo; But it inferred an opposite order between WM and cortex. Other methods did not infer the correct developmental order at all (**Supplementary Fig. S3d**). For mouse coronal hemibrain Stereo-seq data, STAVelo showed similar superiority as applied to other platforms, particularly in inferring the cellular migration direction from Layer 6 to Layer 2/3 (**Fig. 3e**). Other methods failed to infer the correct developmental trend (**Supplementary Fig. S3e**). STAVelo achieved the highest spatial CBDir scores across most cerebral cortex areas and the highest mean CBDir score (**Fig. 3f**). Overall, STAVelo demonstrated highly consistent and accurate results across diverse platforms, reflecting its powerful generalization ability.

### STAVelo infers organ-specific developmental gradients in mouse kidney and chicken heart data

STAVelo revealed critical nephrogenic processes in the mouse kidney Slide-seq data^39^. Following STAGATE-based clustering and spatial domain annotation, both spatial RNA velocity and pseudotime consistently indicated a developmental gradient from proximal convoluted tubule segments (PCT) S1/S2 toward PCT S3, distal convoluted tubules (DCT), and collecting tubes (**Fig. 4a, b**), which is consistent with the published developmental sequence from PCT, DCT to collection tube^40^, but was undetected by other methods (**Supplementary Fig. S4a**). We next identified DKGs by comparing PCT S1/S2 against PCT S3, DCT, and collecting duct compartments. DKGs include SDR, DDR and triple differential rate (TDR) genes, and the Sync pattern accounts for the vast majority of DDR genes (**Supplementary Fig. S4b**). DEGs share less with SDR genes, but more with DDR genes (**Supplementary Fig. S4c**). Representative DKGs, such as *Fnbp4* and *Enah*, are important genes that regulate tubular epithelial migration, luminal budding and segmental elongation, but are not DEGs (**Fig. 4c**). DEGs capture terms associated with post-terminal embryonic maturation and tubular cytoskeleton transporter events. DKGs specifically capture early dynamic regulatory processes such as cell migration, segment boundary formation, and transient phosphorylation signals (**Fig. 4d**).

**Figure 4.**
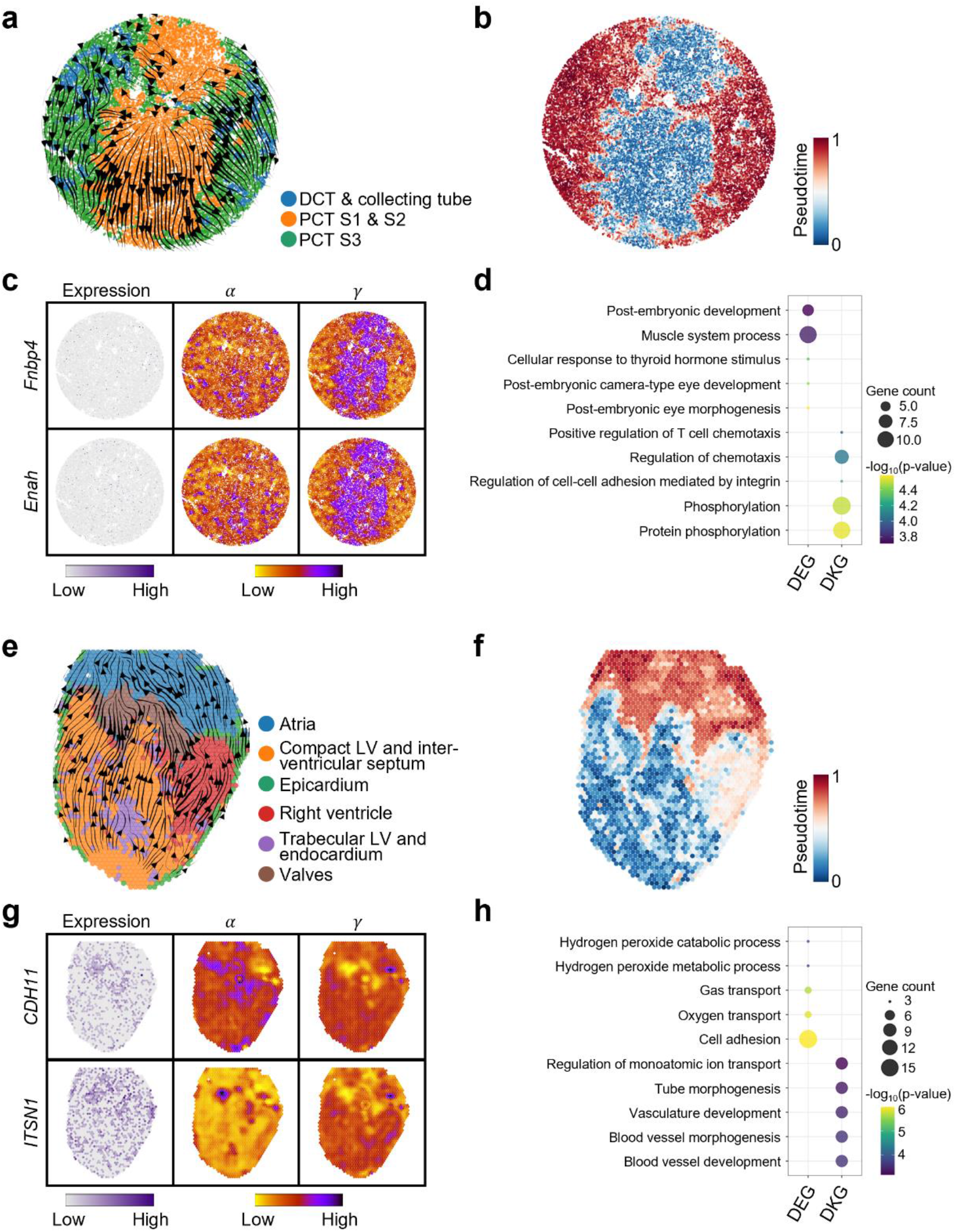
STAVelo infers the important developmental gradient in both the mouse kidney Slide-seq data (a-d) and chicken heart 10x Visium data (e-h). **a** and **e**, Spatial visualization of predicted RNA velocity. Beads are colored by the spatial domains annotated according to the STAGATE clustering results for (**a**), and spots are colored by the spatial domains annotated by the original authors for (**e**). **b** and **f**, Spatial visualization of predicted pseudotime. **c** and **g**, Spatial expression, transcription, and degradation parameter visualizations for representative DKGs. **d** and **h**, Bubble plot showing enriched GO terms of DEGs and DKGs.

We further applied STAVelo to chicken heart 10x Visium data^41^ with spatial domains annotated by the original authors. Spatial RNA velocity and pseudotime reconstructed a cardiogenic axis originating in ventricular myocardium and terminating in atrial chambers (**Fig. 4e, f**), confirming that atrial myocardial differentiation occurs later than ventricular myocardial differentiation^42^. Several other methods did not reflect this trend (**Supplementary Fig. S4d**). We next identified DKGs between ventricular domains (compact LV, interventricular septum, epicardium, RV, trabecular LV and endocardium) and atrial domains (atria and valves). DKGs include SDR and DDR genes, but Async accounts for the most DDR genes (**Supplementary Fig. S4e**). DEGs share fewer genes with either DKGs (**Supplementary Fig. S4f**). Representative DKGs, such as *CDH11* and *ITSN1*, are important genes that regulate inter-myocardial cell adhesion and migration, but are not DEGs (**Fig. 4g**). DEGs only capture functions related to cardiac maturation, such as hydrogen peroxide metabolism and oxygen transport, while DKGs specifically capture functions related to cardiac development, such as ion transport regulation and vessel development (**Fig. 4h**).

In summary, DEGs are more enriched in functions after organ maturation. DKGs are more enriched in functions during organ development, indicating that neural representation of cell-gene-specific kinetic parameters has important biological significance.

### STAVelo reveals spatiotemporal developmental patterns in mouse embryo Stereo-seq data with multiple time points

We applied STAVelo to a mouse embryo Stereo-seq dataset covering five developmental stages from E9.5 to E13.5^43^. Using the published annotations in the original study, we divided cells into existing states, representing cell populations already specified at previous embryonic stages, and newly established states, corresponding to cell populations that acquire distinct, discriminative transcriptional signatures at the current embryonic stages. Across successive developmental stages (E10.5 to E13.5), spatial velocity streams consistently showed directional flows originating from existing states toward newly established ones **(Fig. 5a** and **Supplementary Fig. S5a**). Consistently with spatial observations, complementary UMAP visualizations recapitulated these global developmental trends (**Fig. 5b**).

**Figure 5.**
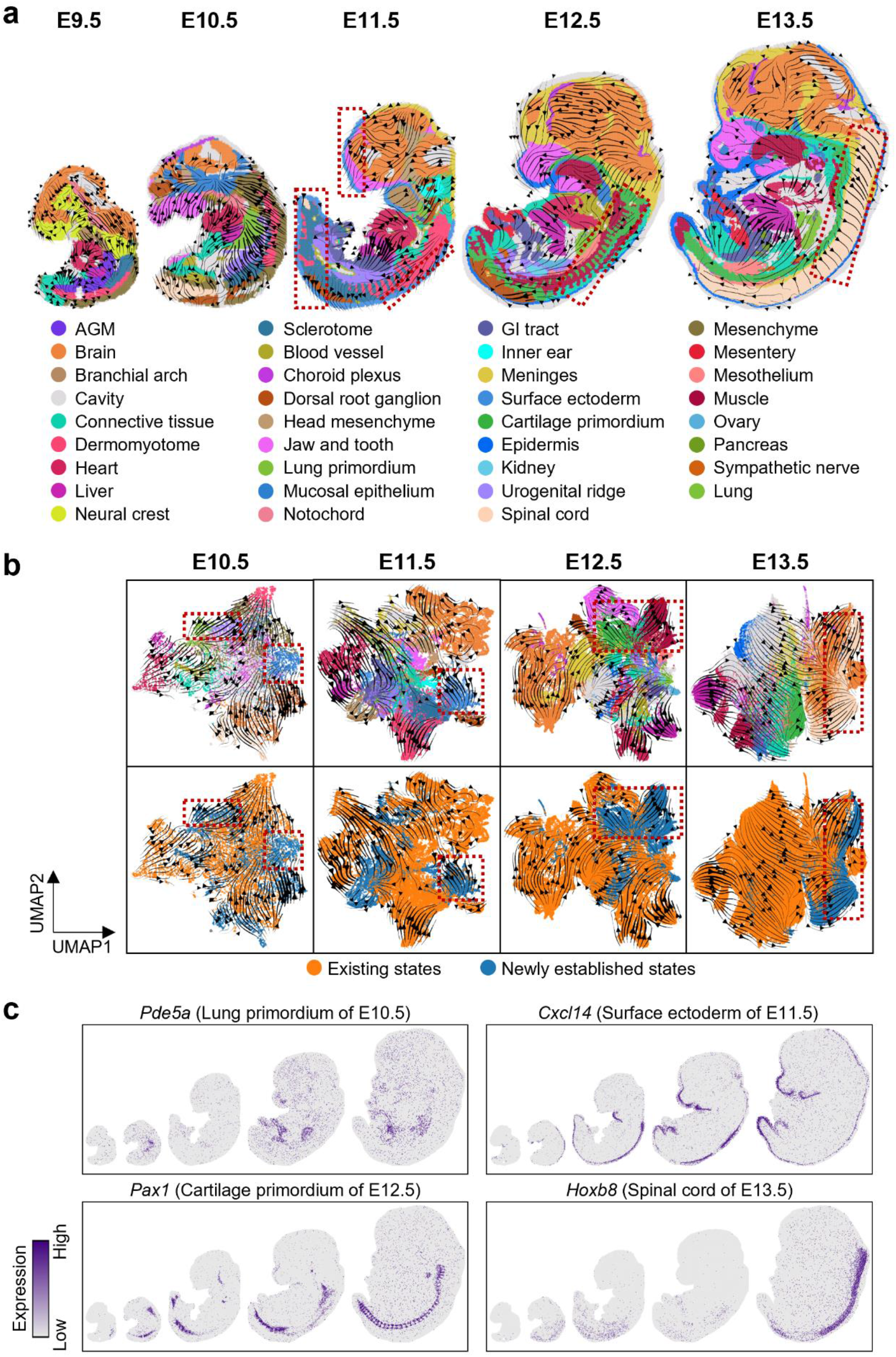
STAVelo revealed spatiotemporal developmental patterns across multiple time points in mouse embryo Stereo-seq data. **a**, Spatial visualization of predicted RNA velocity from E9.5 to E13.5. Cells are colored by the cell types annotated by the original authors. The red dashed boxes represent the newly established states from E10.5 to E12.5 described in the main text. **b**, UMAP visualization of low-dimensional representations with predicted RNA velocity. Cells are colored by the cell types annotated by the original authors in the top panel and development states annotated according to the cell-type annotations in the bottom panel. c, Spatial gene expression of representative marker genes for the newly established states from E10.5 to E12.5 described in the main text.

Representative stage-specific newly established states illustrate this progressive developmental trend. At E10.5, velocity streams target the newly resolvable anatomical primordia, including the lung primordium, urogenital ridge, and mucosal epithelium (**Fig. 5a**). Although their progenitor cell populations already existed at E9.5, these structures acquire distinct transcriptional features and form morphologically distinguishable tissue domains at E10.5. *Pde5a, Kctd14*, and *Penk*, associated with them, were significantly expressed at E10.5 and remained high thereafter (**Fig. 5c** and **Supplementary Fig. S5b, c**). Among them, the lung primordium marker gene *Pde5a* regulates embryonic smooth muscle cell differentiation and later pulmonary vascular tone; It is subsequently highly expressed in the lung, indicating that the lung primordium eventually develops into it. The ovarian granulosa cell marker gene *Kctd14* is initially highly expressed in the urogenital ridge and then in the ovary, consistent with studies on ovarian development. At E11.5, velocity streams target the rapidly developing surface ectoderm (**Fig. 5a**), which will differentiate into the epidermis and can be identified by *Cxcl14* expression, which is associated with the epidermis (**Fig. 5c** and **Supplementary Fig. S5c**). At E12.5, velocity streams target the cartilage primordia that develop from the sclerotome and the muscle that develops from the dermomyotome (**Fig. 5a**), which can be identified by the expression of *Pax1*, which regulates chondrocyte maturation, and *Mymx*, which regulates skeletal muscle development, respectively (**Fig. 5c** and **Supplementary Fig. S5b, c**). At E13.5, velocity streams target the spinal cord (**Fig. 5a**), which already shows significant transcriptional characteristics, as evidenced by *Hoxb8* expression, which plays a key regulatory role in spinal cord development, dorsal neuron differentiation, and sensory conduction pathways (**Fig. 5c** and **Supplementary Fig. S5c**). Collectively, our results resolved the spatiotemporal developmental patterns underlying stepwise organogenesis across E9.5-E13.5 mouse embryogenesis.

### STAVelo uncovers bidirectional migration trajectory from junction zone in mouse placenta Stereo-seq data across multiple developmental stages

To dissect the spatiotemporal cell fate dynamics underlying mouse placenta morphogenesis, we applied STAVelo to the mouse placental Stereo-seq dataset across E7.5 to E12.5^44^. Using the published annotations from the original study, STAVelo illustrated a stage-dependent bidirectional migration trajectory originating from the junctional zone (JZ), based on both spatial observations and complementary UMAP visualizations (**Fig. 6a, b** and **Supplementary Fig. S6a**).

**Figure 6.**
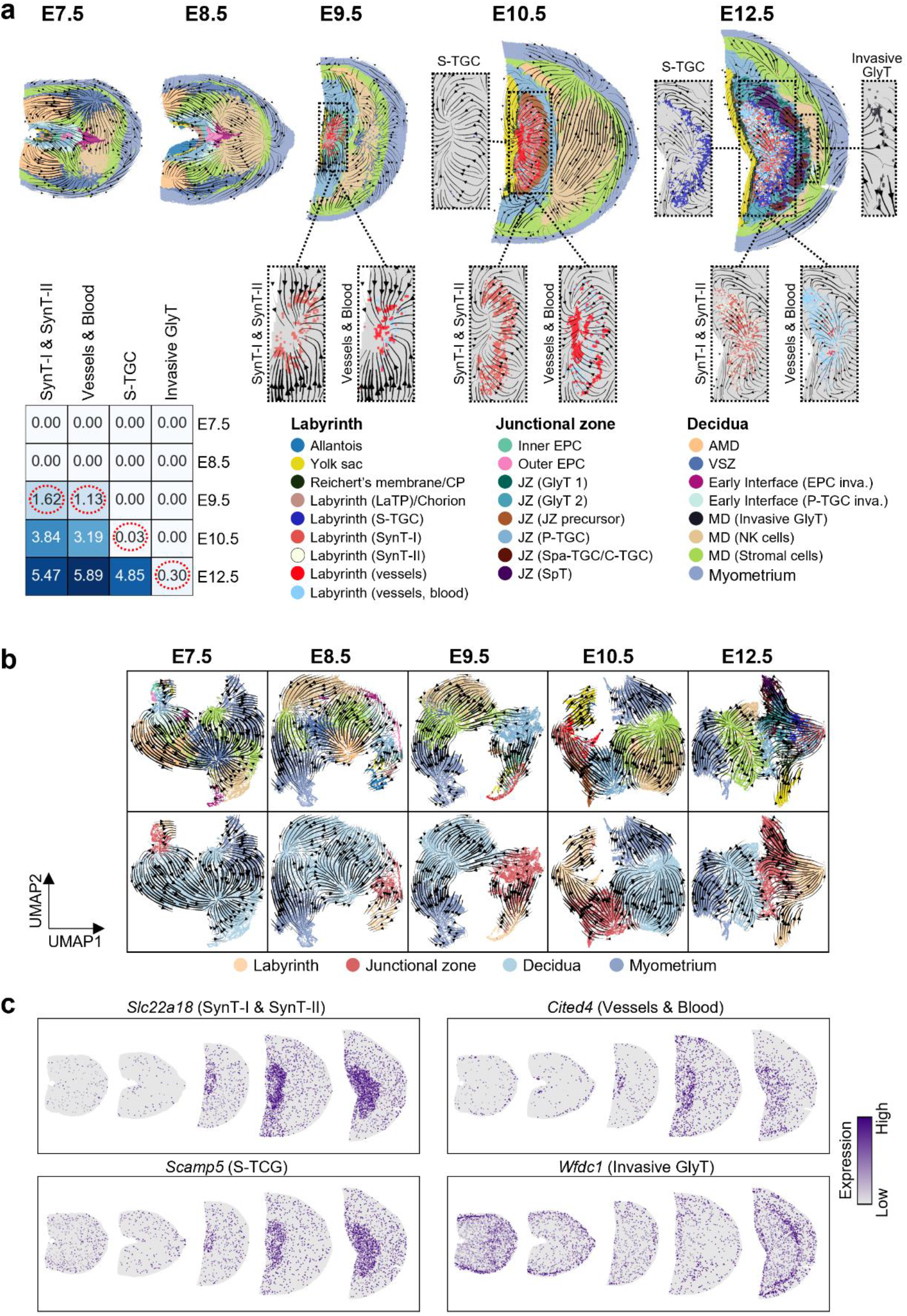
STAVelo revealed bidirectional cell migration patterns across multiple time points in mouse placenta Stereo-seq data. **a**, Spatial visualization of predicted RNA velocity from E7.5 to E12.5. Cells are colored by the cell types annotated by the original authors. The black dashed boxes represent the newly established states from E9.5 to E12.5 described in the main text. Heatmap showing the percentage of the newly established states mentioned in the main text relative to the total number of cells at each time point. SynT, syncytiotrophoblast, S-TGC, sinusoidal trophoblast giant cells, GlyT, glycogen trophoblast. **b**, UMAP visualization of low-dimensional representations with predicted RNA velocity. Cells are colored by the cell types and anatomical regions annotated by the original authors for the top and bottom panels, respectively. c, Spatial gene expression of representative marker genes for the newly established states from E9.5 to E12.5 described in the main text.

At E7.5 and E8.5, the JZ and labyrinth are not yet fully mature, and trophoblast progenitor cells within the JZ gradually differentiate and invasively migrate toward the maternal decidua. From E9.5 to E12.5, the JZ is fully mature and JZ-derived trophoblast progenitor cells migrate to the fetal side and differentiate, constructing the labyrinthine exchange compartments, subsequently producing syncytiotrophoblast (SynT) I/II, blood cells, vessel cells, and sinusoidal trophoblast giant cells (S-TGCs), gradually increasing in proportion as development progresses^44,45^ (**Fig. 6a**). *Cited4, Slc22a18*, and *Scamp5*, as marker genes for the aforementioned cell types, show significantly enhanced spatial signals within the labyrinthine region starting from E9.5 (**Fig. 6c** and **Supplementary Fig. S6b**), reflecting the continuous maturation of structures such as blood vessels and trophoblasts in the fetal labyrinth over developmental stage. Meanwhile, invasive trophoblastic subpopulations, such as glycogen trophoblasts (GlyT), migrate from the JZ to the maternal uterine tissue, remodeling spiral arteries^44,45^ (**Fig. 6a**) and establishing maternal-fetal communication. *Wfdc1*, a GlyT marker gene, was initially detected in the JZ and subsequently spread to the maternal decidua at E12.5 (**Fig. 6c** and **Supplementary Fig. S6b**), reflecting invasive GlyT migration from the JZ to the uterine spiral artery. In short, our spatial velocity results reproduced the bidirectional progenitor cell output of the junctional zone at multiple time points, revealing the cell migration dynamics during mouse placental development.

## Discussion

STAVelo applies a graph attention encoder-decoder neural network to ST data to infer RNA velocity and reconstruct developmental relationships among major tissue structures. STAVelo effectively models spatial information and overcomes the limitation of global uniform kinetic parameters in some methods. To validate the essentiality of spatial modeling, we compared STAVelo with classical non-spatial methods designed for scRNA-seq data. STAVelo correctly predicted relative developmental relationships between tissue structures across multiple species and platforms and showed distinctly better performance than other methods in benchmarking experiments. In particular, STAVelo has also been applied to multi-timepoint mouse embryonic and placental development datasets, revealing their respective spatiotemporal developmental dynamics and demonstrating its application potential on datasets with complex spatiotemporal expression patterns.

STAVelo achieves superior performance on current datasets, yet several extensions remain to be explored. Integrating STAVelo with multi-omics data (e.g., proteomics or metabolomics) could yield deeper insights into transcriptional kinetics. While STAVelo directly infers RNA velocity, it does not simultaneously estimate pseudotime or transcriptional states; developing methods that jointly infer these parameters would enrich analytical outcomes. Furthermore, extending STAVelo to 3D ST datasets would provide a more profound understanding of spatial developmental dynamics. Finally, STAVelo depends on unspliced and spliced expression data, which limits its application to ST datasets without publicly available raw sequencing data. Developing RNA velocity prediction methods on total expression data directly is challenging, but it has broader applications.

STAVelo leverages the graph attention encoder-decoder architecture for data denoising and robust spatial RNA velocity prediction. During model testing, visualization results and quantitative metrics jointly guide network optimization. Rigorous evaluations examined model performance across varying network depths, widths, and attention mechanism configurations. After systematic testing, the finalized network architecture and hyperparameter settings demonstrate exceptional capability in modeling spatial information and predicting spatial RNA velocity, providing a new tool for understanding tissue development at spatial resolution.

## Materials and Methods

### Data preprocessing

#### Upstream processing of the raw sequencing data

For the 10x Visium, Slide-seq, and Slide-seqV2 platforms, we employed Velocyto^15^ to convert the BAM files generated from these platforms to loom files. For the Stereo-seq platform, we employed SAW^46^ to convert the FASTQ files generated from this platform to loom files. These loom files store the unspliced and spliced gene expression counts used for velocity prediction.

#### Preprocessing of gene expression data

We preprocessed the unspliced and spliced counts using the Python package scVelo. First, we performed expression data normalization and highly variable gene selection using the *scvelo*.*pp*.*filter_and_normalize* function. Then we used the *scvelo*.*pp*.*moments* function to compute the top principal components and smooth the expression data. For the ST dataset without spatial domain or cell type annotations, we used STAGATE to cluster spots/beads/cells and annotate them based on specific marker genes.

#### Model implementation

We employed a graph attention encoder-decoder network to model spatial information and predict spatial RNA velocity. Cellular spatial coordinates construct the spatial adjacency graph, while unspliced and spliced expression values serve as node features. The graph and expression data are taken as input to the encoder, where attention mechanisms learn intercellular relationships to generate low-dimensional representations by employing gene expression and spatial information. The decoder outputs transcriptional kinetic rates for RNA velocity estimation. Observed velocities derived from the spatial adjacency graph supervise the neural network training. The trained network predicts RNA velocity vectors mapped to spatial coordinates.

### Spatial adjacency graph construction

Spatial coordinates of cells are used to construct a spatial adjacency graph *G* = (*V, E*). Each vertex *v*_*i*_ represents the gene expression vector *x*_*i*_ = [*u*_*i*_, *s*_*i*_] of cell *i*, where *u*_*i*_ denotes unspliced expression levels and *s*_*i*_ denotes spliced expression levels. An edge *e*_*ij*_ = 1 is established between cells *i* and *j* if their Euclidean distance is below a given threshold.

### Graph attention encoder-decoder network

The graph attention encoder-decoder network learns low-dimensional representations by integrating spatial information and gene expression. This architecture consists of an expression encoder that compresses gene expression data into low-dimensional embeddings, and a kinetics decoder that reconstructs transcriptional kinetic rates from these embeddings.

The model employs attention mechanisms to adaptively learn weights between spatially adjacent cells, reducing the impact of noise and missing data. Standardized gene expression data *X* = [*U, S*] ∈ *R*^*N*×2*D*^ and spatial adjacency matrix *A* ∈ *R*^*N*×*N*^ are fed into the expression encoder, yielding low-dimensional cell representations *Z* in the latent space. The final output comprises transcriptional kinetic rates [α, β, γ] ∈ *R*^*N*×3*D*^, including transcription rate α ∈ *R*^*N*×*D*^, splicing rate β ∈ *R*^*N*×*D*^, and degradation rate γ ∈ *R*^*N*×*D*^. Gene expression serves as the initial cell representation 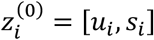.

For an encoder with *L* layers processing *N* cells, the output of cell *i* (*i* ∈ {1,2, …, *N*}) at layer *k* (*k* ∈ {1,2, …, *L* − 1}) is:

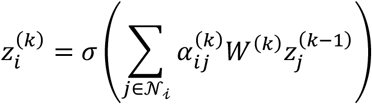

where *N*_*i*_ denotes the neighborhood set of cell *i* (including itself), σ is a nonlinear activation function, *W*^(*k*)^ is the trainable weight matrix at layer *k*, and 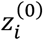 represents initial gene expression *x*_i_. The attention coefficient 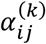 is computed as:

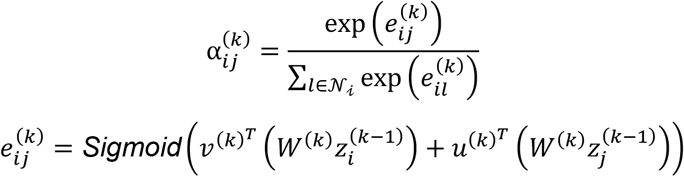

with trainable parameters *v*^(*k*)^ and *u*^(*k*)^. The final encoder layer (*L*) omits attention mechanisms:

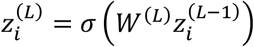

yielding the ultimate low-dimensional representation.

The kinetic decoder employs the same number of layers and reverses the encoding process to reconstruct kinetic rates. Using the expression encoder output as input, intermediate layers (*k* ∈ {2, …, *L* − 1}) reconstruct representations at layer (*k* − 1):

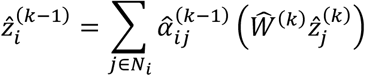

The first decoder layer omits attention:

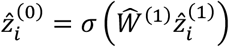

where 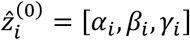 represents kinetic rates. To prevent overfitting, weights are constrained as 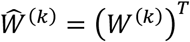 and 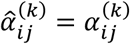.

#### Training procedure

The predicted velocity for each cell is calculated as:

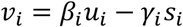

The predicted velocity for unspliced mRNA is given by:

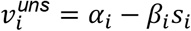

The spatial adjacency graph is utilized to construct observed velocities. Specifically, the observed velocity for cell *i* is derived from expression differences with spatially adjacent cells:

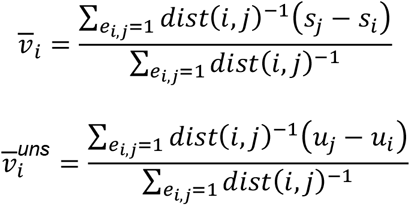

where *dist* denotes the Euclidean distance between cells *j* and *i*.

The training objective maximizes cosine similarity between predicted and observed velocities:

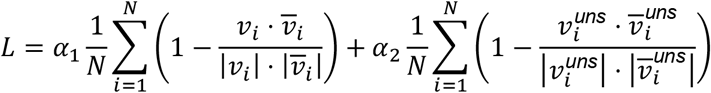

Here, we set α_1_ = α_2_ = 1.

#### Spatial mapping of RNA velocity

To begin, we construct a velocity graph for cells in ST data, where the weight *w*_*ij*_ estimates the magnitude of connectivity. A higher *w*_*ij*_ indicates that the future state of cell *i* is closer to the current state of cell *j*. The weight *w*_*ij*_ is calculated using the cosine similarity between the RNA velocity vector *v*_*i*_ and the gene expression difference *s*_*j*_ − *s*_*i*_:

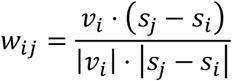

RNA velocity is then projected onto spatial coordinates. Specifically, the transition probability π_*ij*_ from cell *i* to potential target cell *j* is computed using Gaussian-normalized connectivity weights *w*_*ij*_:

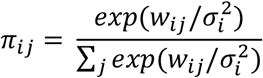

The nuclear width parameter σ_*i*_ for each cell can be locally adjusted between adjacent cells. The spatially mapped RNA velocity *v*_*i*_ is then derived as the weighted sum:

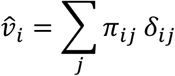

where δ_*ij*_ represents the direction vector pointing from cell *i* to cell *j* in spatial coordinates.

### Pseudotime estimation and differential kinetic gene identification

We estimated pseudotime using the *scvelo*.*tl*.*velocity_pseudotime* function of the Python package scvelo. We specified the root cells in advance based on previous research. We identified the differential kinetic genes (DKGs) using the *scanpy*.*rank_genes_groups* function of the Python package scanpy. Specifically, we set method = *“*t-test*”*, and layer = *“*alpha*”, “*beta*”*, and *“*gamma*”* for differential analysis of transcription, splicing, and degradation parameters, respectively. For each kinetic parameter, we considered genes with adjusted *p*-value < 0.05 and absolute log_2_ fold change > 1 to be significantly different for that parameter. We identified the genes with at least one significantly different kinetic parameter as DKGs. We further classified the DKGs with one, two, and three significantly different kinetic parameters as single differential rate (SDR), double differential rate (DDR), and triple differential rate (TDR). We classified the DDR genes into the synchronous (Sync) and asynchronous (Async) patterns. We identified the DDR genes with an upregulated transcription rate and a downregulated degradation rate, or a downregulated transcription rate and an upregulated degradation rate, as the Sync pattern, and the others as the Async pattern.

### Biological process enrichment analysis

We performed GO enrichment analysis on identified DEGs and DKGs using the *enrichGO* function of the R package clusterProfiler^47^. We considered the pathways with *p*-values < 0.05 to be significantly enriched.

### Benchmarking methods

#### Velocyto

Velocyto assumes cells reside in stable equilibrium where expression profiles exhibit no net changes over time, implemented via the *scvelo*.*tl*.*velocity()* function with parameter mode=*‘*deterministic*’*.

#### scVelo

In contrast to Velocyto, scVelo extends this framework by incorporating temporal dimensions of cellular kinetics to provide comprehensive views of cell trajectories while capturing complex interactions between cellular states, considering both temporal gene expression variations and underlying data graph structures for velocity estimation. scVelo explicitly accounts for cellular transition directionality and potential multi-steady-state existence, requiring sequential execution of *scvelo*.*tl*.*recover_dynamics()* followed by *scvelo*.*tl*.*velocity()* with mode=*‘*dynamical*’*.

#### cellDancer

cellDancer employs fully connected neural networks to model relationships between gene expression and kinetic rates. The RNA velocity is computed via *celldancer*.*velocity()* and *celldancer*.*compute_cell_velocity()* functions.

#### Evaluation Metrics

To provide a comprehensive evaluation, we adopt two metrics commonly used in RNA velocity estimation for scRNA-seq data to assess spatial velocity. This approach assumes ST data has been partitioned into distinct clusters via manual annotation or clustering, where each cluster represents a specific cell type or tissue structure.

#### Cross-boundary direction correctness score

Spatial cross-boundary direction correctness (CBDir) score evaluates the directional accuracy of RNA velocities between clusters, assuming biologically supported developmental trajectories from cluster A to B. Let *C*_*A*_ and *C*_*B*_ be cells of clusters A and B, and *N*(*i*) be the neighboring cells of cell *i*. The CBDir score for cell *i* is calculated as:

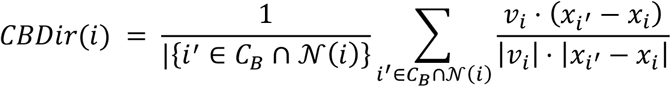

where 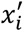 and *x*_*i*_ represent spatial coordinates of cells *i* and *i’*, and *v*_*i*_ denotes the low-dimensional RNA velocity vector at spatial location *i*.

#### In-cluster coherence score

Spatial in-cluster velocity coherence (ICVCoh) score assesses velocity consistency within a single cluster via cosine similarity. The ICVCoh score for cell *i* is defined as:

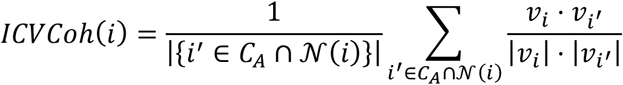

Unlike CBDir, ICVCoh evaluates velocity coherence but cannot assess directional correctness.

## Data availability

All datasets used in this study (**Supplementary Table S1**) are publicly available and can be accessed through the following sources: The human DLPFC Visium data: https://interactive.libd.org/spatialLIBD (gene expression data), https://research.libd.org/globus (BAM files); The mouse brain Visium data: https://10xgenomics.com/datasets/adult-mouse-brain-coronal-section-visium-ff-1-standard (gene expression data and BAM files of coronal section), https://10xgenomics.com/datasets/mouse-brain-serial-section-1-sagittal-posterior-1-standard-1-1-0 (gene expression data and BAM files of sagittal posterior section); The mouse hippocampus Slide-seqV2 data: Single Cell Portal (SCP) database under the accession code SCP815 (gene expression data and BAM files); The mouse cortex Slide-seqV2 data: SCP database under the accession code SCP2082 (gene expression data and BAM files); The mouse coronal hemibrain Stereo-seq data: https://www.dropbox.com/scl/fi/8raqrh3kchb81qubahe6g/mousebrain_bin60_clustered.h5ad?rlkey=0jivz6n8xbhnhuxd1u90sy7ri&e=1 (processed h5ad file with unspliced and spliced data); The mouse kidney Slide-seq data: SCP database under the accession code SCP354 (gene expression data and BAM files); The chicken heart Visium data: Gene Expression Omnibus (GEO) database under the accession code GSE149457 (gene expression data), https://zenodo.org/records/6798659 (processed h5ad file with unspliced and spliced data); The mouse embryo Stereo-seq data: https://db.cngb.org/stomics/mosta (gene expression data), https://db.cngb.org/search/project/CNP0001543 (FASTQ files); The mouse placentation Stereo-seq data: https://db.cngb.org/stomics/mpsta/download/ (processed h5ad file with unspliced and spliced data).

## Code availability

The STAVelo algorithm is implemented in Python and is available upon request.

## Acknowledgments

This work has been supported by the Postdoctoral Fellowship Program of CPSF (no. GZC20252011), the Fundamental Research Funds for the Central Universities, the National Natural Science Foundation of China (nos. 32341013, 12326614 to S.Z), and the CAS Project for Young Scientists in Basic Research (no. YSBR-034 to S.Z.).

## Author contributions

Shihua Zhang conceived and supervised the project. Shang Li designed and implemented STAVelo algorithm. Shang Li and Qunlun Shen validated the study. Shang Li, Qunlun Shen and Shihua Zhang wrote the manuscript. All authors read and approved the final manuscript.

## Competing interests

The authors declare no competing interests.

## Notes

### Competing Interest Statement

The authors have declared no competing interest.

