## Supplemental Figures for "STAVelo: deciphering spatiotemporal cellular dynamics via spatial RNA velocity inference from spatial transcriptomics data"

### Supplementary Figures

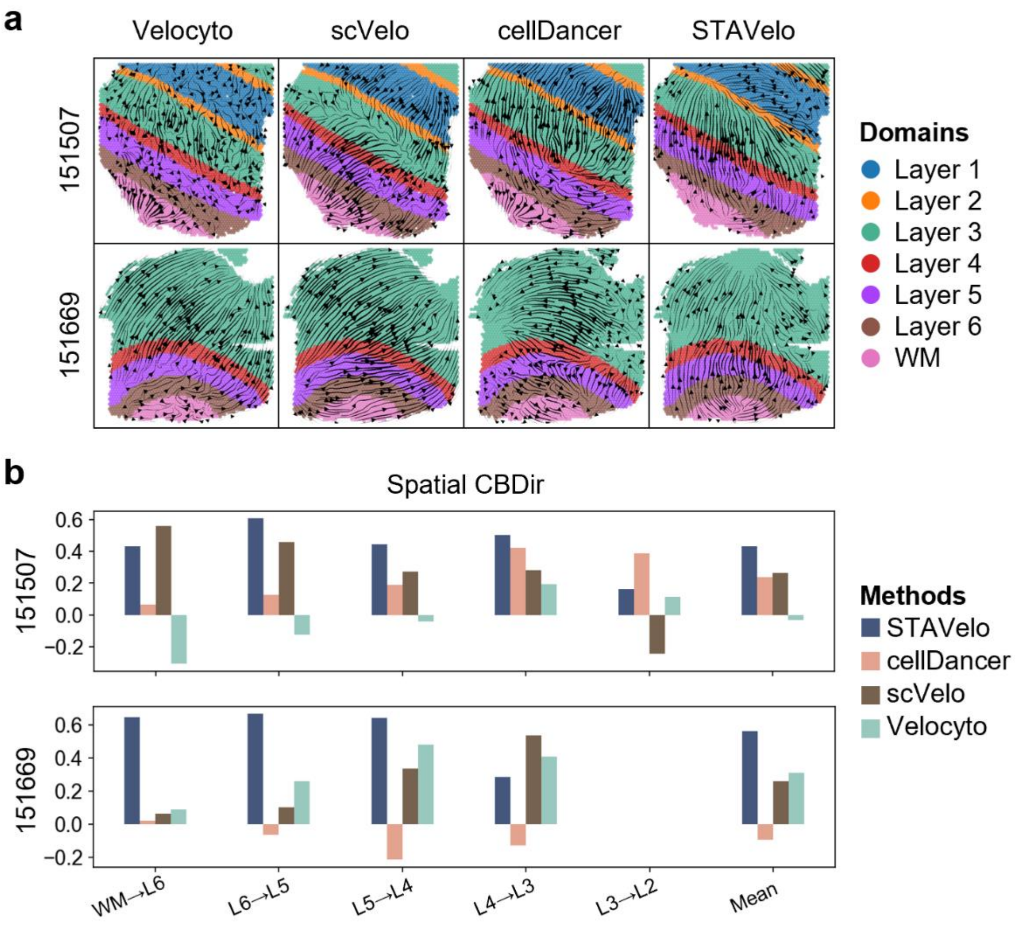


**Figure S1. Spatial RNA velocity inferred by STAVelo and competing methods on the human DLPFC sections 151508 and 151669. a,** Spatial visualization of RNA velocity predicted by STAVelo, cellDancer, scVelo, and Velocyto. **b,** Bar chart showing spatial CBDir scores of STAVelo, cellDancer, scVelo, and Velocyto.

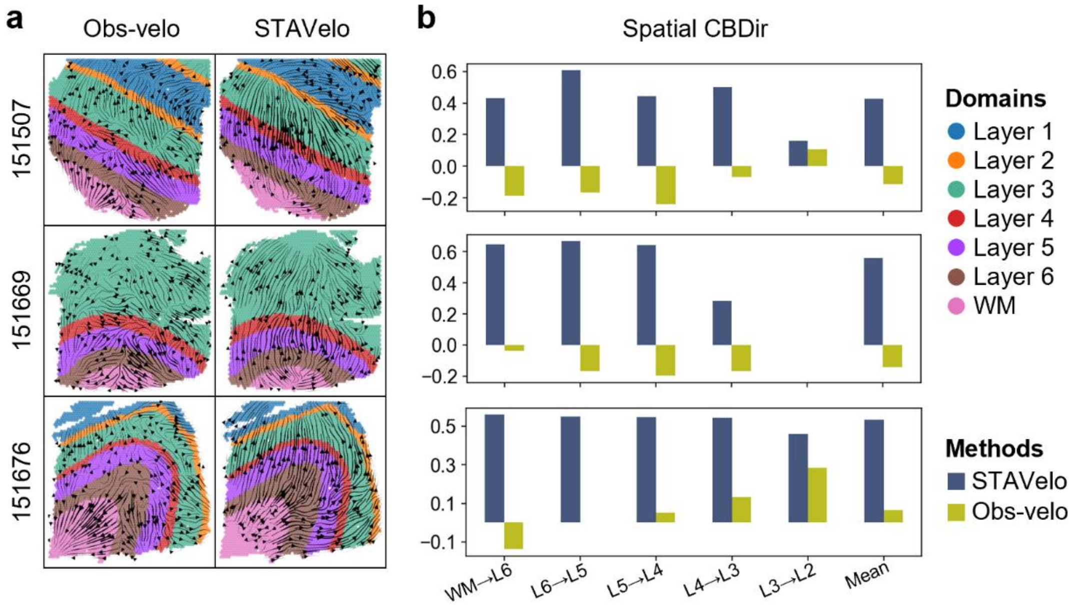


**Figure S2. Ablation experiments for the graph attention encoder-decoder network of STAVelo on the human DLPFC sections 151508, 151669, and 151676. a,** Spatial visualization of RNA velocity predicted by STAVelo and Obs-velo. **b,** Bar chart showing spatial CBDir scores of STAVelo and Obs-velo.

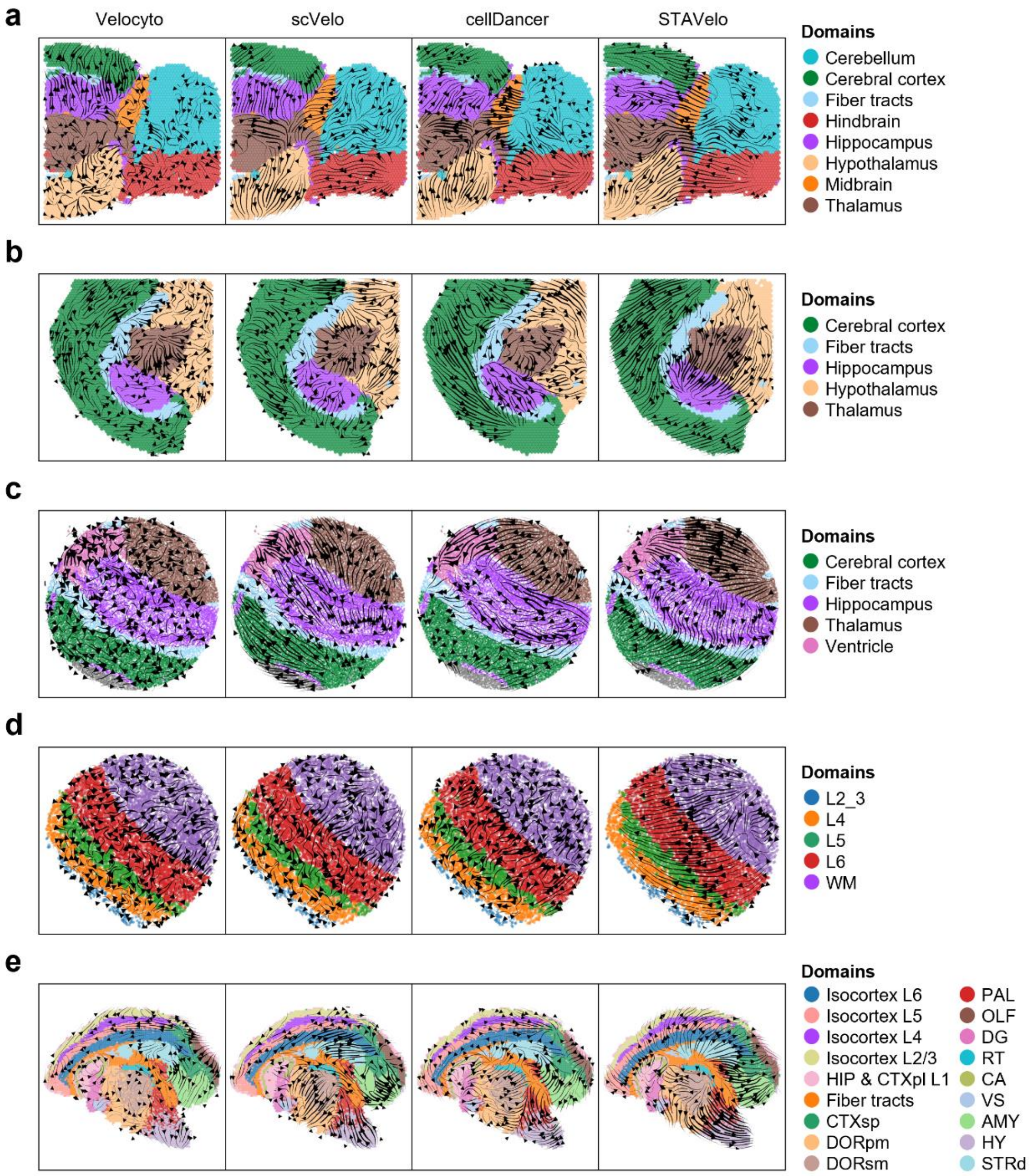


**Figure S3. Spatial visualization of RNA velocity inferred by STAVelo, cellDancer, scVelo, and Velocyto on the mouse brain data.** These data refers to different sections and platforms including, 10x Visium posterior-sagittal section (**a**), 10x Visium coronal section (**b**), Slide-seq coronal section (**c**), Slide-seq cortex section (**d**), and Stereo-seq hemibrain coronal section (**e**).

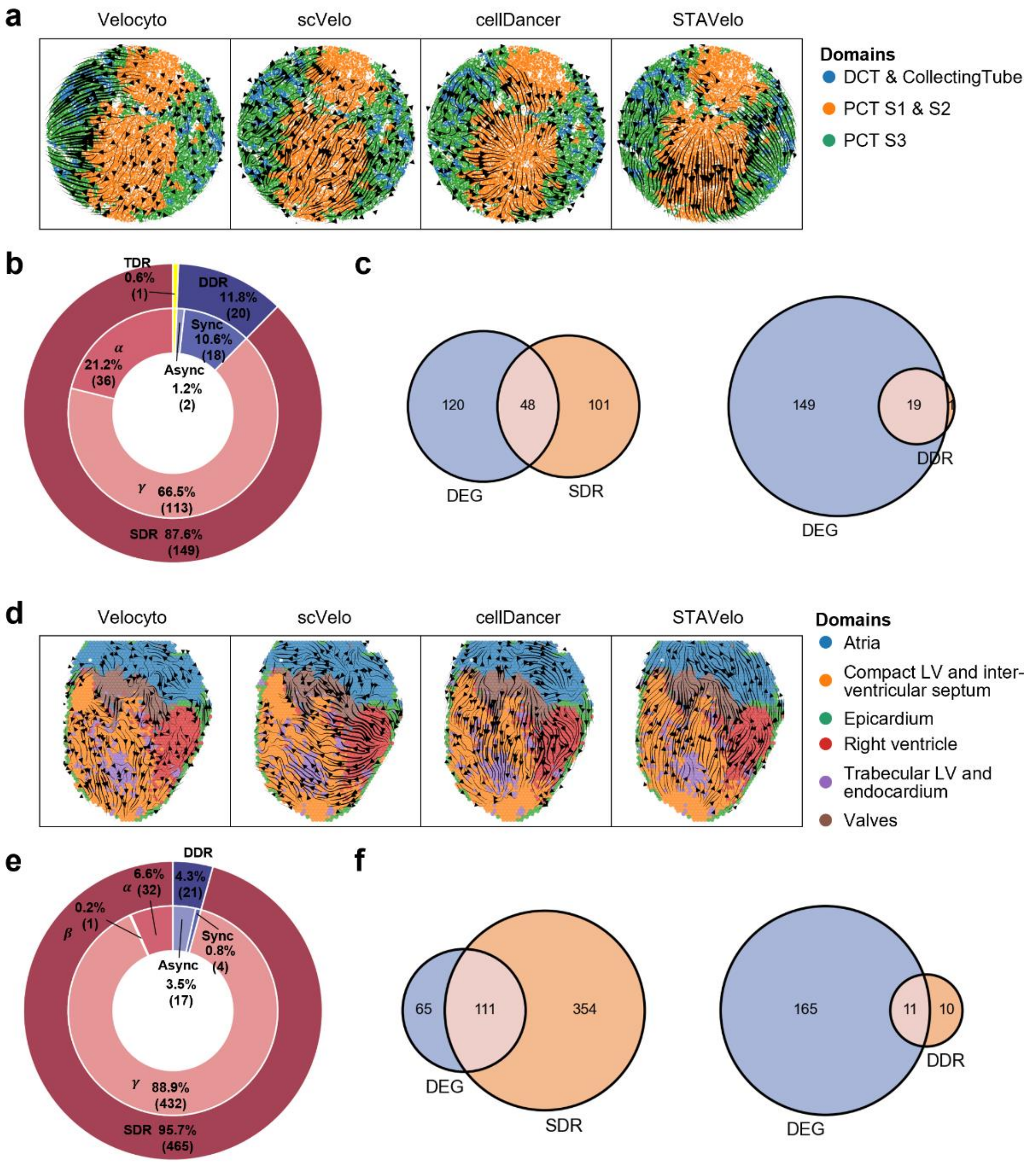


**Figure S4. Spatial RNA velocity inferred by STAVelo and competing methods on the mouse kidney Slide-seq data (a-c) and chicken heart 10x Visium data (d-f). a** and **d,** Spatial visualization of RNA velocity predicted by STAVelo, cellDancer, scVelo, and Velocyto. **b** and **e,** Double doughnut chart showing the proportion of each category of differential kinetic genes (DKGs), including single differential rate (SDR), double differential rate (DDR) and triple differential rate (TDR) categories, with DDR further divided into synchronous (Sync) and asynchronous (Async) patterns. **c** and **f,** Venn diagrams illustrating overlap between differential expression genes (DEGs), SDR and DDR genes.

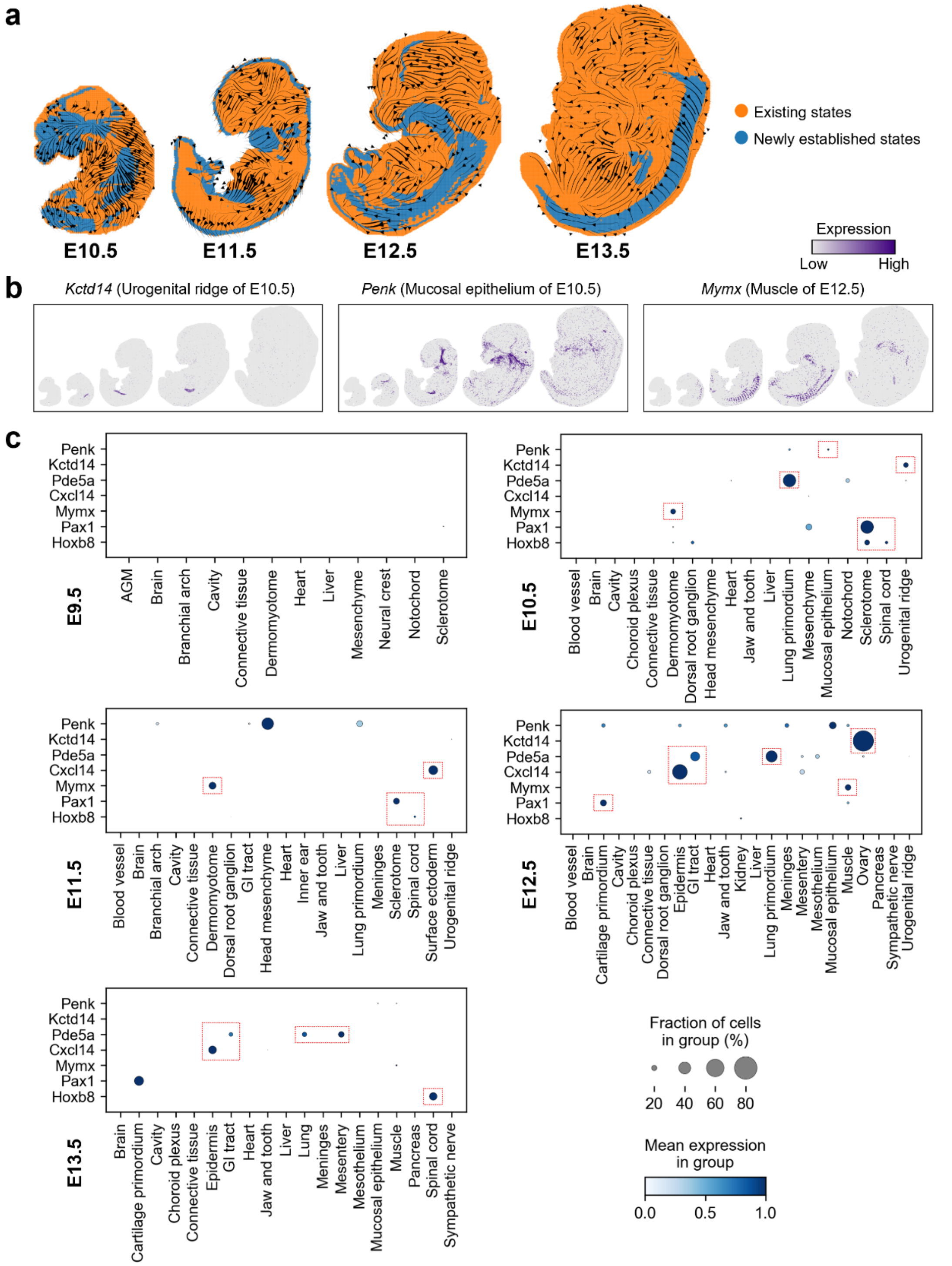


**Figure S5. Spatial RNA velocity inferred by STAVelo on the mouse embryo Stereo-seq data across multiple timepoints. a,** Spatial visualization of predicted RNA velocity from E9.5 to E13.5. Cells are colored by the development states annotated according to the cell types annotations. **b**, Spatial gene expression of representative marker genes for the newly established states from E10.5 and E12.5 described in the main text. **c,** Dotplot showing the expression of representative marker genes across the cell types at each time point.

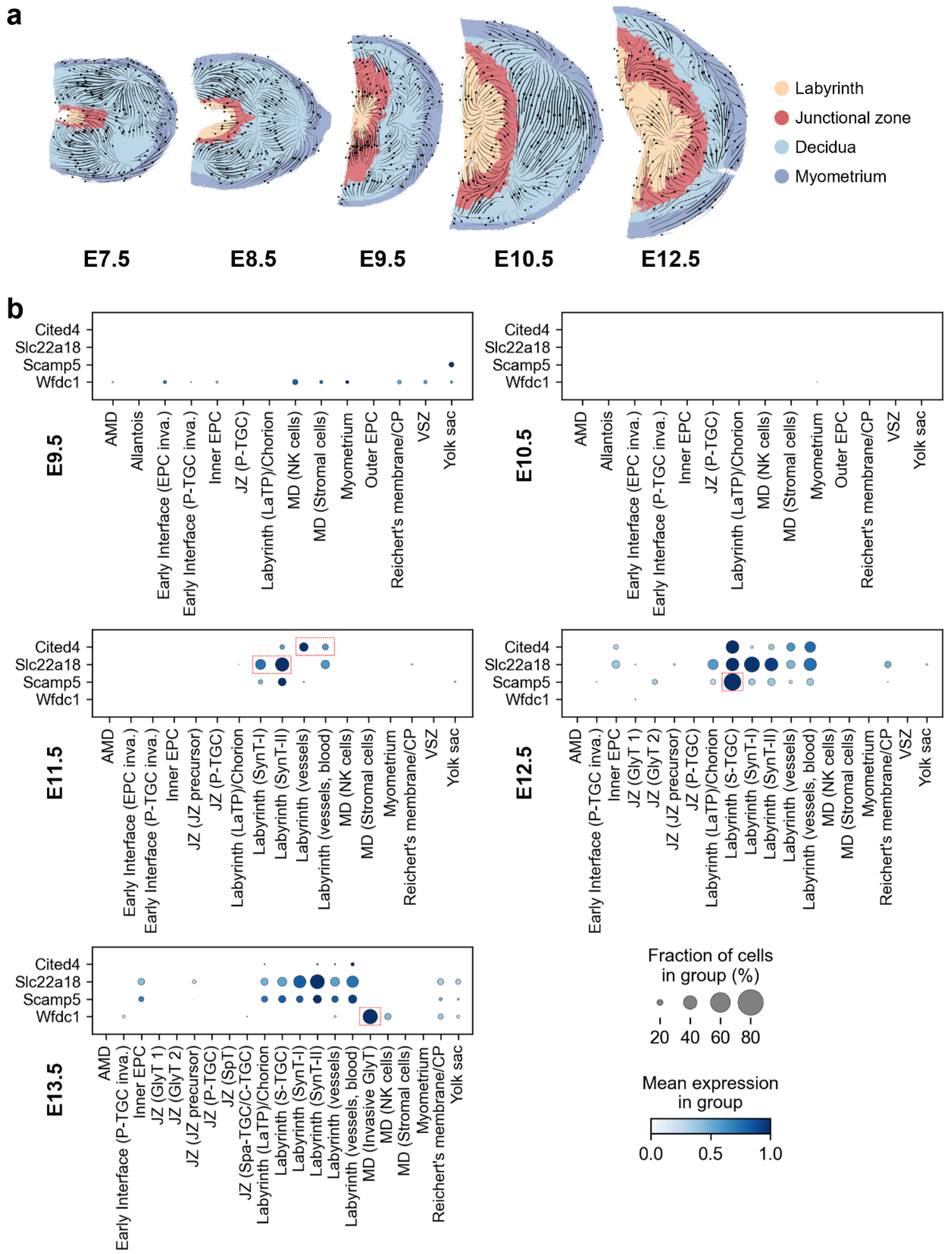


**Figure S6. Spatial RNA velocity inferred by STAVelo on the mouse placenta Stereo-seq data across multiple timepoints. a,** Spatial visualization of predicted RNA velocity from E7.5 to E12.5. Cells are colored by the anatomical regions annotated by the original authors. **b,** Dotplot showing the expression of representative marker genes across the cell types at each time point.

**Supplementary Table**

**Table S1. Description of all ST datasets used in this study.**

| Platform | Tissue | Section | Related figures | Reference |
|---|---|---|---|---|
| **10x Visium** | Human dorsolateral prefrontal cortex (DLPFC) | 151508, 151669, 151676. | Fig. 2, Fig. S1, Fig. S2 | (1) |
| | Mouse brain | Coronal section | Fig. 3, Fig. S3 | 10x Visium demo |
| | | Sagittal-posterior section | Fig. 3, Fig. S3 | |
| | Chicken heart | Day 14 | Fig. 4, Fig. S4 | (2) |
| **Slide-seq** | Mouse brain | Puck_200115_08 | Fig. 3, Fig. S3 | (3) |
| | Mouse brain | Puck_210824_01 | Fig. 3, Fig. S3 | (4) |
| | Mouse kidney | Puck_180528_23 | Fig. 4, Fig. S4 | (5) |
| **Stereo-seq** | Mouse brain | Coronal hemibrain section | Fig. 3, Fig. S3 | (6) |
| | Mouse embryo | E9.5, E10.5, E11.5, E12.5, E13.5 | Fig. 5, Fig. S5 | (7) |
| | Mouse placenta | E7.5, E8.5, E9.5, E10.5, E12.5, E14.5 | Fig. 6, Fig. S6 | (8) |
